# Structure-function analysis of the nsp14 N7-guanine methyltransferase reveals an essential role in *Betacoronavirus* replication

**DOI:** 10.1101/2021.05.17.444407

**Authors:** Natacha S. Ogando, Priscila El Kazzi, Jessika C. Zevenhoven-Dobbe, Brenda W. Bontes, Alice Decombe, Clara C. Posthuma, Volker Thiel, Bruno Canard, François Ferron, Etienne Decroly, Eric J. Snijder

## Abstract

As coronaviruses (CoVs) replicate in the host cell cytoplasm, they rely on their own capping machinery to ensure the efficient translation of their mRNAs, protect them from degradation by cellular 5’ exoribonucleases, and escape innate immune sensing. The CoV nonstructural protein 14 (nsp14) is a bi-functional replicase subunit harboring an N-terminal 3′-to-5′ exoribonuclease (ExoN) domain and a C-terminal (N7-guanine)-methyltransferase (N7-MTase) domain that is presumably involved in viral mRNA capping. Here, we aimed to integrate structural, biochemical, and virological data to assess the importance of conserved N7-MTase residues for nsp14’s enzymatic activities and virus viability. We revisited the crystal structure of severe acute respiratory syndrome (SARS)-CoV nsp14 to perform an *in silico* comparative analysis between betacoronaviruses. We identified several residues likely involved in the formation of the N7-MTase catalytic pocket, which presents a fold distinct from the Rossmann fold observed in most known MTases. Next, for SARS-CoV and Middle East respiratory syndrome-CoV, site-directed mutagenesis of selected residues was used to assess their importance for *in vitro* enzymatic activity. Most of the engineered mutations abolished N7-MTase activity, while not affecting nsp14-ExoN activity. Upon reverse engineering of these mutations into different betacoronavirus genomes, we identified two substitutions (R310A and F426A in SARS-CoV nsp14) abrogating virus viability and one mutation (H424A) yielding a crippled phenotype across all viruses tested. Our results identify the N7-MTase as a critical enzyme for betacoronavirus replication and define key residues of its catalytic pocket that can be targeted to design inhibitors with a potential *pan*-coronaviral activity spectrum.

**Significance Statement:** The ongoing SARS-CoV-2 pandemic emphasizes the urgent need to develop efficient broad-spectrum anti-CoV drugs. The structure-function characterization of conserved CoV replicative enzymes is key to identifying the most suitable drug targets. Using a multidisciplinary comparative approach and different betacoronaviruses, we characterized the key conserved residues of the nsp14 (N7-guanine)-methyltransferase, a poorly defined subunit of the CoV mRNA-synthesizing machinery. Our study highlights the unique structural features of this enzyme and establishes its essential role in betacoronavirus replication, while identifying two residues that are critical for the replication of the four betacoronaviruses tested, including SARS-CoV-2.

## Introduction

At their 5′ end, all eukaryotic mRNAs carry an N7-methylguanosine cap that ensures their translation by mediating mRNA recognition during the formation of the ribosomal pre-initiation complex. The co-transcriptional capping of cellular pre-mRNAs occurs in the nucleus and is also critical for pre-mRNA splicing and nuclear export (reviewed in (1–3)). The mRNA cap consists of an N7-methylated 5’ guanosine moiety that is linked to the first nucleotide of the transcript by a 5’-5’ triphosphate bridge (4). Its synthesis requires (presumably) the consecutive involvement of triphosphatase, guanylyltransferase, and guanine-N7 methyltransferase activities to produce a cap-0 structure. The first nucleotides of mammalian mRNAs are then methylated on the 2’OH position to yield a cap-1 structure that identifies the transcript as “self” and prevents activation of innate immune sensors (reviewed in (2, 5)). Furthermore, the cap structure promotes mRNA stability by providing protection from cellular 5’ exoribonucleases.

Viruses rely on host ribosomes for their gene expression and have adopted different strategies to ensure translation of their own mRNAs. These include using the canonical nuclear capping pathway, so-called ‘cap-snatching’ mechanisms, and replacement of the cap by a ribosome-recruiting RNA structure (reviewed in (2, 6, 7)). Various cytosolically replicating virus families have evolved their own capping machinery. The latter applies to the coronavirus (CoV) family, which includes the severe acute respiratory syndrome coronavirus 2 (SARS-CoV-2), the causative agent of COVID-19 (8, 9), and a range of other CoVs infecting human or animal hosts (10, 11). This century alone, the CoV family has given rise to three major zoonotic introductions: SARS-CoV-2, the Middle East respiratory syndrome-CoV (MERS-CoV) discovered in 2012, and SARS-CoV, emerging in South East Asia in 2002. All three belong to the genus *Betacoronavirus*, which is abundantly represented among CoVs circulating in bat species (12–15). Despite their demonstrated potential to cross species barriers, prophylactic and therapeutic solutions for CoV infections to prevent or rapidly contain the current COVID-19 pandemic were not available.

The positive-sense CoV genome is unusually large (∼30 kb) and its 5’ proximal two-thirds encodes for two replicase polyproteins that are post-translationally cleaved into 16 nonstructural proteins (nsp) (16, 17). The CoV replicative enzymes, including the nsp12 RNA-dependent RNA polymerase (RdRp), assemble into a protein complex that is embedded within virus-induced replication organelles (18–20) and directs the synthesis and capping of newly made viral genomes as well as subgenomic mRNAs that serve to express additional CoV genes. Capping is thought to involve the successive action of multiple CoV enzymes: (i) the nsp13 RNA triphosphatase removing the γ phosphate from the nascent 5’-triphosphorylated RNA (21, 22); (ii) an RNA guanylyltransferase (GTAse) producing a GpppN cap by transferring guanosine monophosphate (GMP) to the RNA’s dephosphorylated 5’ end, a role recently attributed to the nsp12 nucleotidyltransferase (NiRAN) domain, but remaining to be confirmed (23–25); (iii) the nsp14 (N7-guanine)-methyltransferase (N7-MTase) methylating the N7 position of the cap while using S-adenosyl methionine (SAM) as methyl donor; (iv) the nsp16 ribose 2’-O-methyltransferase (2’-O-MTase) converting the cap-0 into a cap-1 structure (^7m^GpppN_2’Om_; (26, 27)) by performing additional methylation with the assistance of nsp10 as co-factor (26, 28, 29).

Over the past 15 years, the CoV capping machinery has mainly been analyzed *in vitro*, in particular for SARS-CoV, but its characterization in the context of the viral replication cycle has remained limited to a handful of studies. This applies in particular to the CoV N7-MTase domain, expressed as part of the ∼60-kDa nsp14, a bi-functional replicase subunit also containing an N-terminal 3’-to-5’ exoribonuclease domain implicated in promoting the fidelity of CoV replication (30, 31). Following the discovery of an N7-MTase activity associated with nsp14’s C-terminal domain (27), the protein was found to methylate non-methylated cap analogues or guanosine triphosphate (GTP) substrates in the presence of SAM in biochemical assays (26, 32, 33). While the association of nsp10 with nsp14 enhances its ExoN activity, the *in vitro* N7-MTase activity does not depend on nsp10 co-factor (26, 34). Biochemical and structural characterization of the N7-MTase and ExoN domains demonstrated that the two domains are functionally distinct (35–38). Nevertheless, truncations and alanine substitutions in the ExoN domain can severely affect SAM binding and N7-MTase activity (27, 33). The notion that the two enzymatic domains are structurally intertwined was also supported by the SARS-CoV nsp14 crystal structure (35, 36) which was found to be composed of (i) a flexible N-terminal sub-domain forming the nsp10 binding site (aa 1-58), (ii) the 3’-to-5’ exoribonuclease (ExoN) domain (aa 1-291), (iii) a flexible hinge region consisting of a loop that connects the N- and C-terminal domains, and three strands protruding from the C-terminal domain (aa 285-300 and aa 407-430), and (iv) the C-terminal N7-MTase domain (aa 292-527) ((35, 36); Fig. 1A).

**Figure 1.**
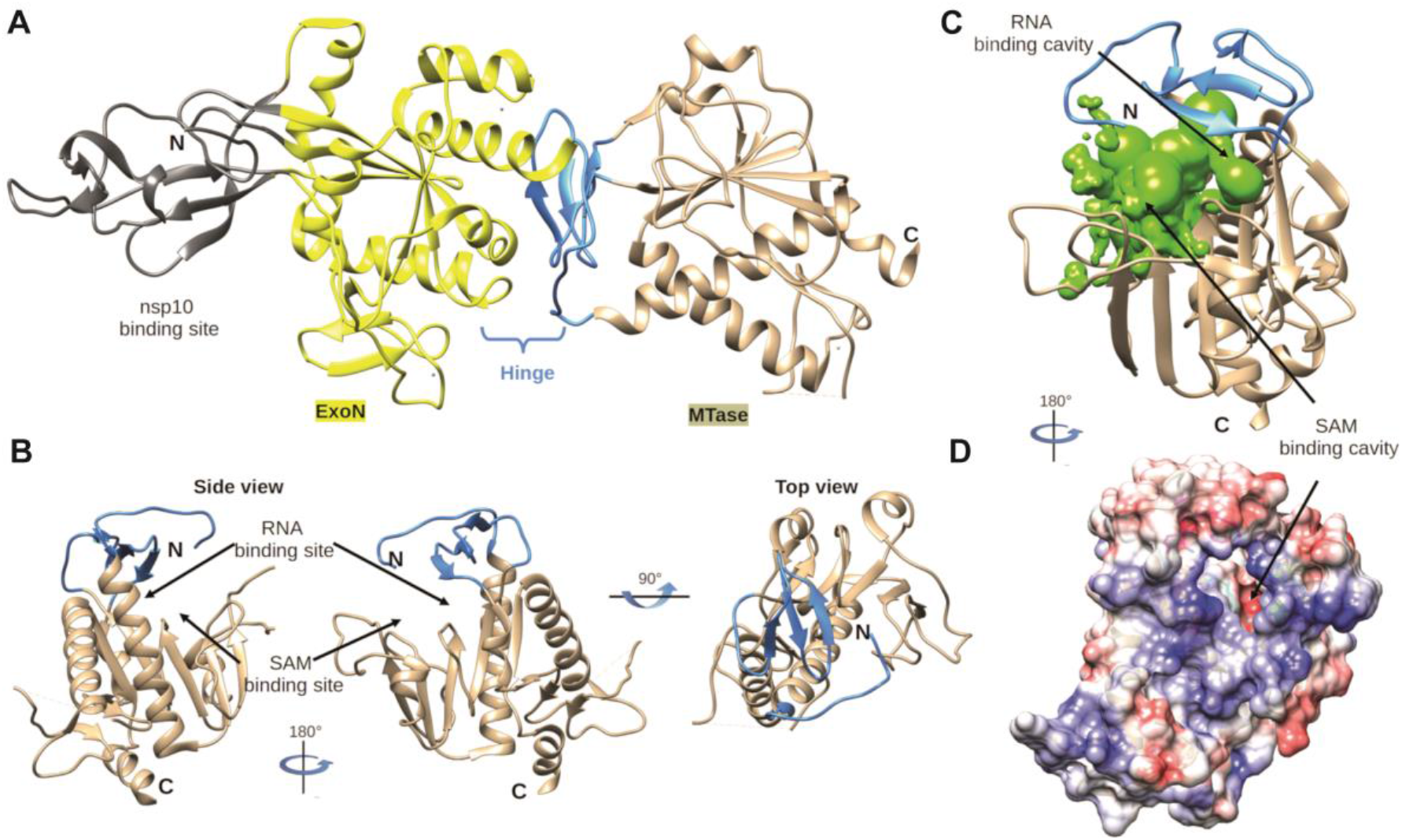
Global architecture of coronavirus nsp14. (A) Architecture of SARS-CoV nsp14 (PDB 5NFY) showing the nsp10 binding site (grey), N-terminal ExoN domain (yellow), hinge subdomain (blue), and C-terminal N7-MTase domain (brown). (B) Side and top view of the hinge region and N7-MTase domain. The three strands of the hinge (blue) protrude from the N7-MTase domain (brown). (C) Analysis of the volume of the N7-MTase active site, with the cavity highlighted in green and hinge subdomain in blue. (D) Electrostatic surface representation of the CoV nsp14 hinge region and N7-MTase domain. Surface electrostatic potential calculated by Adaptive Poisson-Boltzmann Solver, from - 10 (red) to + 10 (blue) kT/e.

Interestingly, the structural analysis of the SARS-CoV-nsp14 N7-MTase revealed a non-Rossmann fold (36), distinguishing this enzyme from commonly known cellular and viral methyltransferases (39, 40). Despite the biochemical characterization of the CoV N7-MTase, the assessment of its importance for virus replication has remained limited to studies with a few point mutations introduced into nsp14 of murine hepatitis virus, a model betacoronavirus (41–44). These studies highlighted two motifs important for CoV replication: (i) the presumed SAM binding motif I (DxGxPxG/A, with x being any amino acid; Fig. 2C, motif III), first discovered by superimposition of a SARS-CoV nsp14 N7-MTase structure model with the crystal structures of cellular N7-MTases (27); (ii) nsp14 residues 420-428 (Fig. 2C, part of motif VI) that, based on the SARS-CoV crystal structure, seem to form a constricted pocket holding the cap’s GTP moiety (35). Comparative analysis of N7-MTase domains revealed that a number of residues crucial for substrate and ligand binding are conserved among homologous enzymes in more distant CoV (Fig. 2A) and other nidoviruses (45–47).

**Figure 2.**
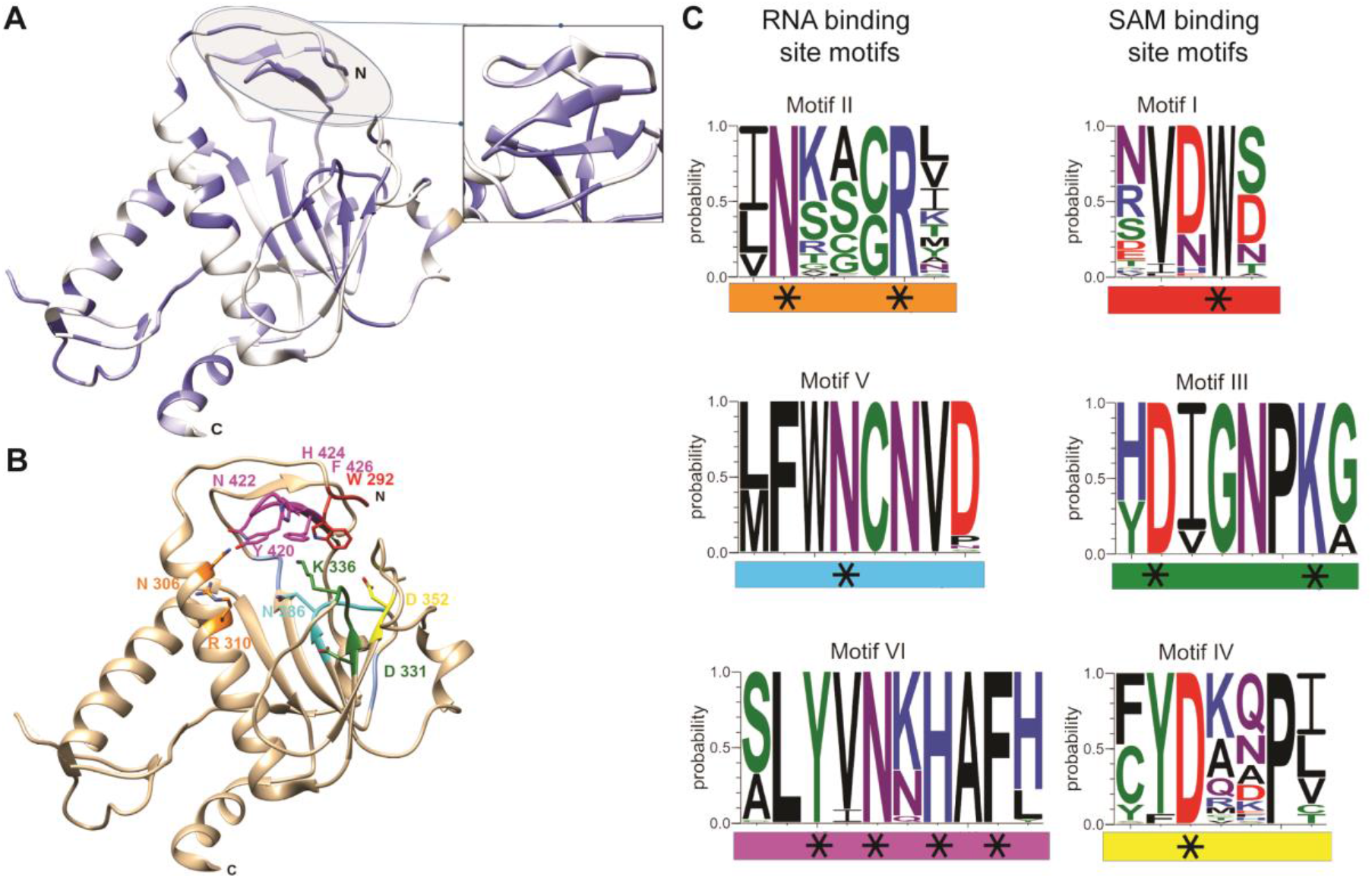
Coronavirus-wide nsp14 N7-MTase conservation and structural analysis. (A) CoV nsp14 amino acid sequence conservation plotted on the structure (PDB 5NFY) of the SARS-CoV hinge region and N7-MTase domain (dark blue to white shading representing 100% to less than 50% sequence identity). A list of sequences used for this comparison is presented in Table S1. (B) Close-up of identified conserved motifs and residues in the N7-MTase catalytic pocket. (C) WebLogo representation of 6 conserved motifs (I-VI) identified in the N7-MTase catalytic pocket. Each motif is highlighted with a specific color (matching that in panel B) and categorized as a proposed SAM- or RNA-binding motif. Black stars highlight charged or aromatic residues most likely involved in ligand binding or catalytic activity.

Due to its conservation and unique structural features, the CoV N7-MTase constitutes an attractive target for antiviral drug development (48–50), to combat SARS-CoV-2 or future emerging CoV threats. Only a few compounds have been reported to inhibit nsp14 N7-MTase activity *in vitro* (26, 32, 48–50). Evaluation of their antiviral activity revealed limited inhibition of CoV replication in cell culture, suggesting poor bio-availability and/or specificity (48, 51). Structural, biochemical, and virological studies of CoV N7-MTase structure and function have not been integrated thus far. Here, we set out to define the catalytic pocket, characterize its involvement in enzymatic activity, and use these observations to probe the enzyme’s importance for CoV replication. Using four different betacoronaviruses (SARS-CoV, MERS-CoV, MHV, and SARS-CoV-2), we identified conserved features and residues supporting N7-MTase activity and viral replication, thus providing a solid framework for future efforts to design broad-spectrum inhibitors of this critical CoV enzyme.

## Results

### Identification of key residues for RNA and SAM binding by the CoV N7-Mtase

The previously resolved SARS-CoV nsp14 structure (35, 36) revealed how the ExoN and N7-MTase domains are structurally interconnected, with possible functional implications (Fig. 1). Thus far, a structure of nsp14 in complex with 5’-capped RNA is lacking. Due to some structural peculiarities, it was unclear which conserved residues may be mechanistically involved in N7-methylation and how important these may be for overall CoV replication. Therefore, we first revisited the core structure of the SARS-CoV N7-MTase, to guide a subsequent biochemical and virological comparison across multiple betacoronaviruses.

In the SARS-CoV nsp14 structure (35), the ExoN core presents a fold characteristic of the DED/EDh family of exonucleases (31, 52, 53). However, the N7-MTase domain does not exhibit the canonical ‘Rossmann fold’ that is common among RNA virus MTases, RNA cap-0 MTases at large, and all five classes of SAM-dependent MTases (54, 55). A hinge region that is highly conserved across CoVs is present at the interface of nsp14’s ExoN and N7-MTase domains (Fig. 1A) and constitutes a unique structural feature of this bi-functional CoV protein. It not only connects the two domains, but also forms an extension that protrudes from the surface of the N7-MTase domain (Fig. 1B). Although, the overall structure suggests ExoN and N7-MTase to be separate domains, the successful expression and purification of truncated forms of the N7-MTase domain, with or without the hinge sub-domain, has not been reported (27, 56). This might be related to the hydrophobic nature of the hinge, which is likely important for protein stability and folding. Several studies reported that the replacement of ExoN catalytic residues does not impair the N7-Mtase activity, suggesting that the functional interplay between the two domains is limited (26, 27, 33, 37, 38, 48). Whereas the hinge region allows lateral and rotational movement of the two nsp14 domains, one side of the hinge also constitutes the ‘ceiling’ of the N7-MTase active site (Fig. 1B).

The structures of SARS-CoV nsp14 in complex with SAM and GpppA (PDB: 5C8S and 5C8T; (35)) have defined the enzyme’s cap-binding pocket. However, the crystal packing profoundly constrained the structural characterization of the N7-MTase domain and the overall low resolution left uncertainties regarding the positioning of the RNA ligand. Therefore, we performed a thorough structural analysis of the enzyme’s cavity, supported by CoV-wide nsp14 sequence comparisons, in order to define conserved N7-MTase residues that may be involved in enzymatic activity (Fig. S2). Several aspects were taken into consideration while delimiting the SAM and RNA binding sites: the general geometry of the cavity, its electrostatic properties, and the conservation of specific amino acid residues. We used Surfnet software (57) to define the volume corresponding to the ligand-binding cavity (Fig. 1C). This volume is shaped as a dual bulb, with the larger pocket accommodating the capped RNA and the smaller one forming the SAM binding site. An electrostatic surface analysis shows positive charges lining the wall of the putative RNA-binding cavity (Fig. 1D and Fig. S1), which would be consistent with its function. Likewise, positive charges that might accommodate the carbocyclic part of the methyl donor were identified in the SAM binding pocket (Fig. 1D). Additionally, conserved hydrophobic residues (Motif I; Fig. 2C) were mapped to a deep hydrophobic cavity, supposedly accommodating the SAM base by a stacking interaction with F426 (SARS-CoV numbering). Finally, the integration of the structural models with CoV-wide N7-MTase sequence comparisons (Table S1 and Fig. S2) allowed the identification of conserved potential key residues within each cavity (blue regions in Fig. 2A). Based on their conservation and positioning, six conserved motifs (I-VI) were defined, each containing a series of specific charged or aromatic residues that have their side chain pointing toward the cavity (Fig. 2B and 2C). Their features suggested they can facilitate the methyl transfer from SAM onto the cap’s guanine residue at the 5’ end of the RNA substrate, by stabilizing and/or correctly positioning the cap structure. The following potential key residues were identified (amino acid numbers matching those in SARS-CoV nsp14): Motif I, W292; Motif II, N306 and R310; Motif III, D331 and K336; Motif IV, D352; Motif V, N386; Motif VI, Y420, N422, H424, and F426 (Fig. 2B and 2C). To assess the possible impact of their replacement on nsp14 folding, we analyzed the predicted impact of single-site substitutions with alanine on the thermostability of SARS-CoV nsp14 (Table S2 and Fig. 5A). Except for R310, all replacements yielded positive ΔΔG values, suggesting that these mutations may affect MTase stability by altering either its fold, or the cavity for SAM or RNA binding (Table S2 and Fig. 5A). Noticeably, mutations in Motifs I and VI, which are spatially close as part of the hinge and most likely involved in the binding of capped RNA, resulted in the largest ΔΔG gains. Similar observations were made when the impact of substitutions with other amino acids was evaluated for other betacoronaviruses (Table S3).

### Identification of residues crucial for *in vitro* N7-MTase activity

To experimentally verify the outcome of our structural analysis (Fig. 1–2), we probed the functional importance of selected residues through targeted mutagenesis and *in vitro* N7-MTase assays. Based on their conservation, charge, position, and potential role for RNA or SAM binding in the catalytic pocket (Fig. 2B and 2C), eleven and nine N7-MTase residues were replaced with alanine in recombinant SARS-CoV and MERS-CoV nsp14, respectively. N-terminally H-tagged proteins were expressed in *E*. coli and purified using immobilized metal affinity chromatography (IMAC) followed by size exclusion chromatography (Fig. 3A and 3B).

**Figure 3.**
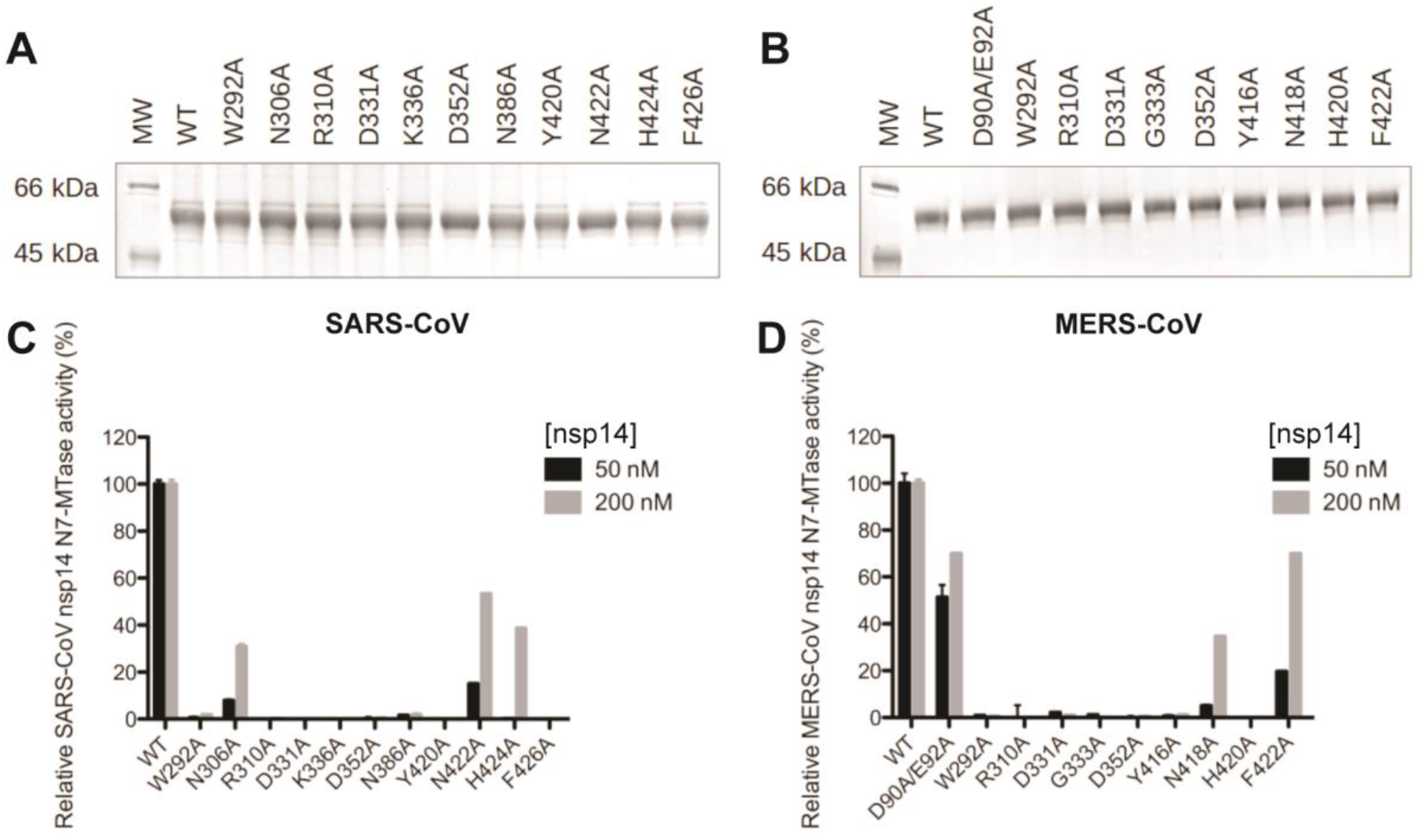
Expression and *in vitro* N7-MTase activity of SARS-CoV and MERS-CoV nsp14 mutants. Recombinant SARS-CoV (A) and MERS-CoV (B) wild-type and mutant nsp14 proteins were expressed in *E. coli* and purified. Proteins were loaded (2 µg and 1 µg for SARS-CoV and MERS-CoV, respectively) and analyzed using 10% SDS-PAGE gels stained with Coomassie blue. The *in vitro* N7-MTase activity of SARS-CoV (C) and MERS-CoV (D) nsp14 mutants was determined using an assay with a GpppACCCC synthetic RNA substrate and radiolabeled SAM as methyl donor. Nsp14 concentrations of 50 and 200 nM were used, as indicated. N7-MTase activities were compared to those of the respective wild-type nsp14 controls. For MERS-CoV, ExoN knockout mutant D90A/E92A was included as a control.

We evaluated the N7-MTase activity of nsp14 mutants in an assay using a GpppACCCC capped RNA substrate and radiolabeled [^3^H]SAM. The transfer of the [^3^H]methyl group onto the RNA substrate was quantified using filter binding assays (Fig. 3C and 3D), as described previously (26, 34), and compared to the enzymatic activity of wild-type SARS-CoV or MERS-CoV nsp14. With the exception of N306A (30% residual activity), N422A (53% residual activity), and H424A (40% remaining), all SARS-CoV mutations tested almost completely abrogated nsp14 N7-MTase activity (Fig. 3C). In the case of MERS-CoV nsp14, only mutants N418A and F422A retained partial N7-MTase activity, 34% and 70%, respectively, while again all other mutations rendered the enzymatic activity barely detectable (Fig. 3D). In terms of residual activity, differences were observed for some pairs of equivalent SARS-CoV and MERS-CoV mutants (e.g. the H and F in motif VI), but overall the results were fully in line with the outcome of our structural analysis. Thus, our data confirmed and extended a previous study (35), and showed that N7-MTase activity is affected by mutations that either may inhibit SAM binding (W292A, D331A, G333A, K336A, D352A in SARS-CoV) or likely interfere with RNA chain stabilization (N306A, R310A, Y420A, N422A, F426A) in the catalytic pocket.

### Revisiting the interplay between the N7-MTase and ExoN domains of nsp14

Despite the notion that the ExoN and N7-MTase domains of CoV nsp14 may be functionally independent (27, 33, 35, 36), they are structurally interconnected by the hinge region (Fig. 1). Therefore, we evaluated the impact of all of our N7-MTase mutations on ExoN functionality, using an *in vitro* assay with 5’-radiolabeled RNA substrate H4 (34), a 22-nt RNA of which the largest part folds into a hairpin structure. Its degradation was monitored using denaturing polyacrylamide gel electrophoresis and autoradiography (Fig. 4). Nsp10 was added as a co-factor that importantly stimulates nsp14 ExoN activity (34, 35, 37), as again confirmed in the ‘nsp14 only’ control assay (Fig. 4). As expected, in time course experiments, we observed the progressive 3’-to-5’ degradation of the RNA substrate by the wild-type nsp10-nsp14 pair of both SARS-CoV (Fig. 4A) and MERS-CoV (Fig. 4B). In the same assay, most of our N7-MTase mutations barely affected ExoN activity (Fig. 4A and 4B), also supporting the notion that these mutant proteins had folded correctly. In contrast, the ExoN activity of SARS-CoV mutants R310A, Y420A and H424A, and MERS-CoV mutant W292A was strongly or partially affected, as indicated by the reduced amount of hydrolysis products at the bottom of the gel. Meanwhile, incorporation of the H420A mutation completely abrogated MERS-CoV nsp14 ExoN activity. Three of the five mutations (Y420A and H424A in SARS-CoV and H420A in MERS-CoV) that affected ExoN activity mapped to motif VI in the hinge region (Fig. 2). Based on the structural analysis, we assume that these mutations affect either the overall nsp14 folding or – more likely - constrain the flexibility of the hinge subdomain with negative consequences for ExoN functionality (35, 36). Conversely, a MERS-CoV ExoN knockout mutant (D90A/E92A), which was included as a control, was found to modestly impact N7-MTase activity (Fig. 3D). Taken together, our data suggest that, although the N7-MTase sequence is well conserved among betacoronaviruses ((35, 37) and Fig. S2), the differences observed between SARS-CoV and MERS-CoV must be caused by a certain level of structural variability or differences in recombinant protein stability.

**Figure 4.**
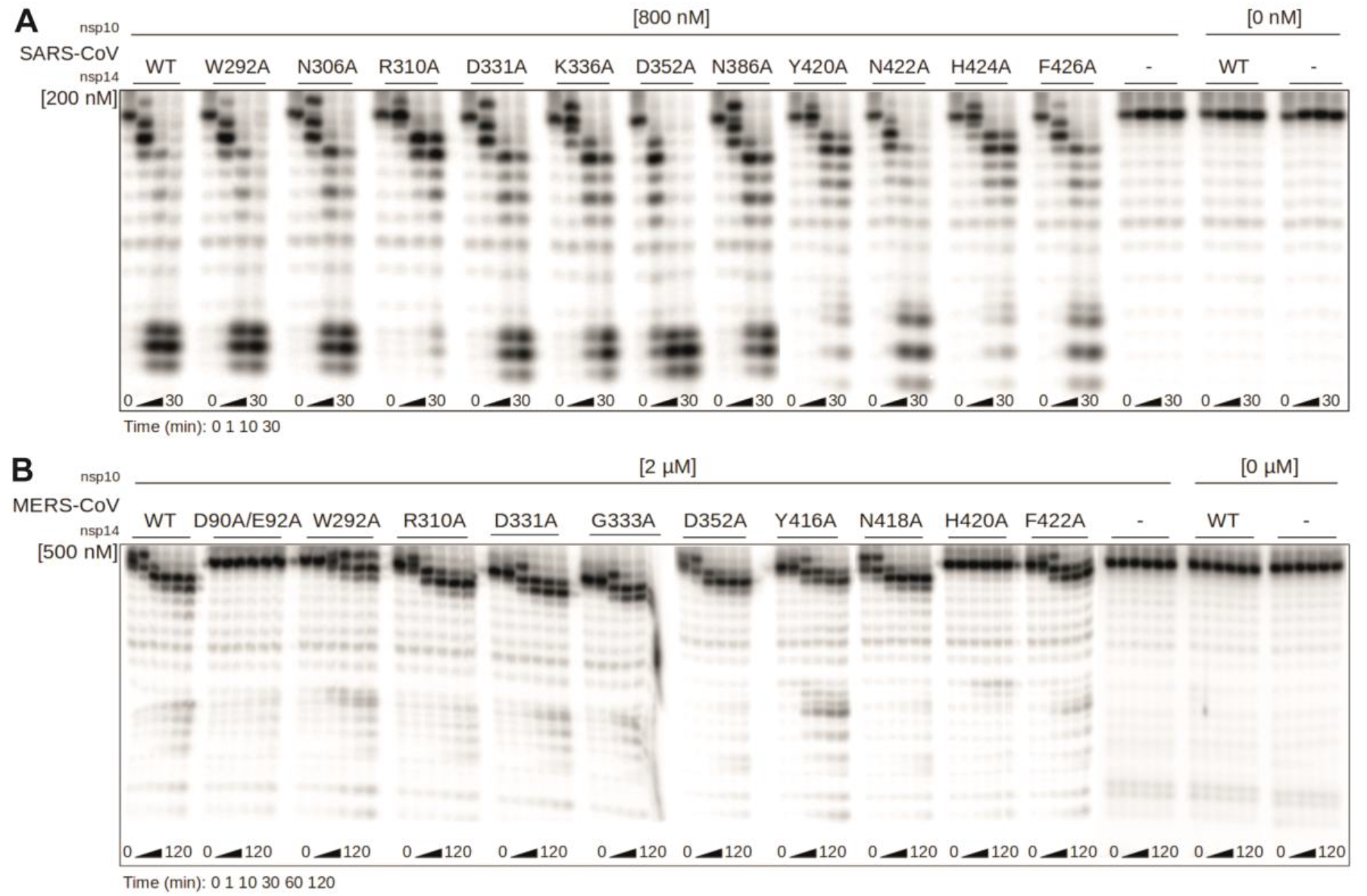
*In vitro* exoribonuclease activity of SARS-CoV and MERS-CoV N7-MTase mutants. The *in vitro* ExoN activity of SARS-CoV (A) and MERS-CoV (B) mutant nsp14 proteins (Fig. 3) was determined by monitoring the degradation of a 5’ radiolabeled RNA substrate (see Methods). An nsp14 concentration of 200 or 500 nM was used (as indicated) and a fourfold molar excess of the corresponding nsp10 was added. A time course assay was performed using time points 0, 1, 10, and 30 min for SARS-CoV, and 0, 1, 10, 30, 60, and 120 min for MERS-CoV nsp14. Reaction products were analyzed by denaturing gel electrophoresis and autoradiography.

### The nsp14 N7-MTase is critical for SARS-CoV viability

As summarized above, most prior biochemical and structural studies of the CoV N7-MTase were performed using SARS-CoV nsp14, whereas mutagenesis in the context of virus replication (using reverse genetics) was restricted to MHV studies in which, for different reasons, the conserved D and G residues in motif III and the Y residue in motif VI were targeted (41, 43, 58). To establish a connection between the biochemical and virological data on the N7-MTase, we first introduced twelve single N7-MTase mutations into the SARS-CoV genome, using a bacterial artificial chromosome-based reverse genetics system. Each mutant was engineered in duplicate and launched by *in vitro* transcribing full-length RNA that was electroporated into BHK-21 cells. To propagate viral progeny, if released, transfected BHK-21 cells were mixed with Vero E6 cells and incubated up to 6 days. Each mutant was launched at least four times, using RNA from 2 independent clones in two independent experiments, and mutant phenotypes are summarized in Fig. 5A.

**Figure 5.**
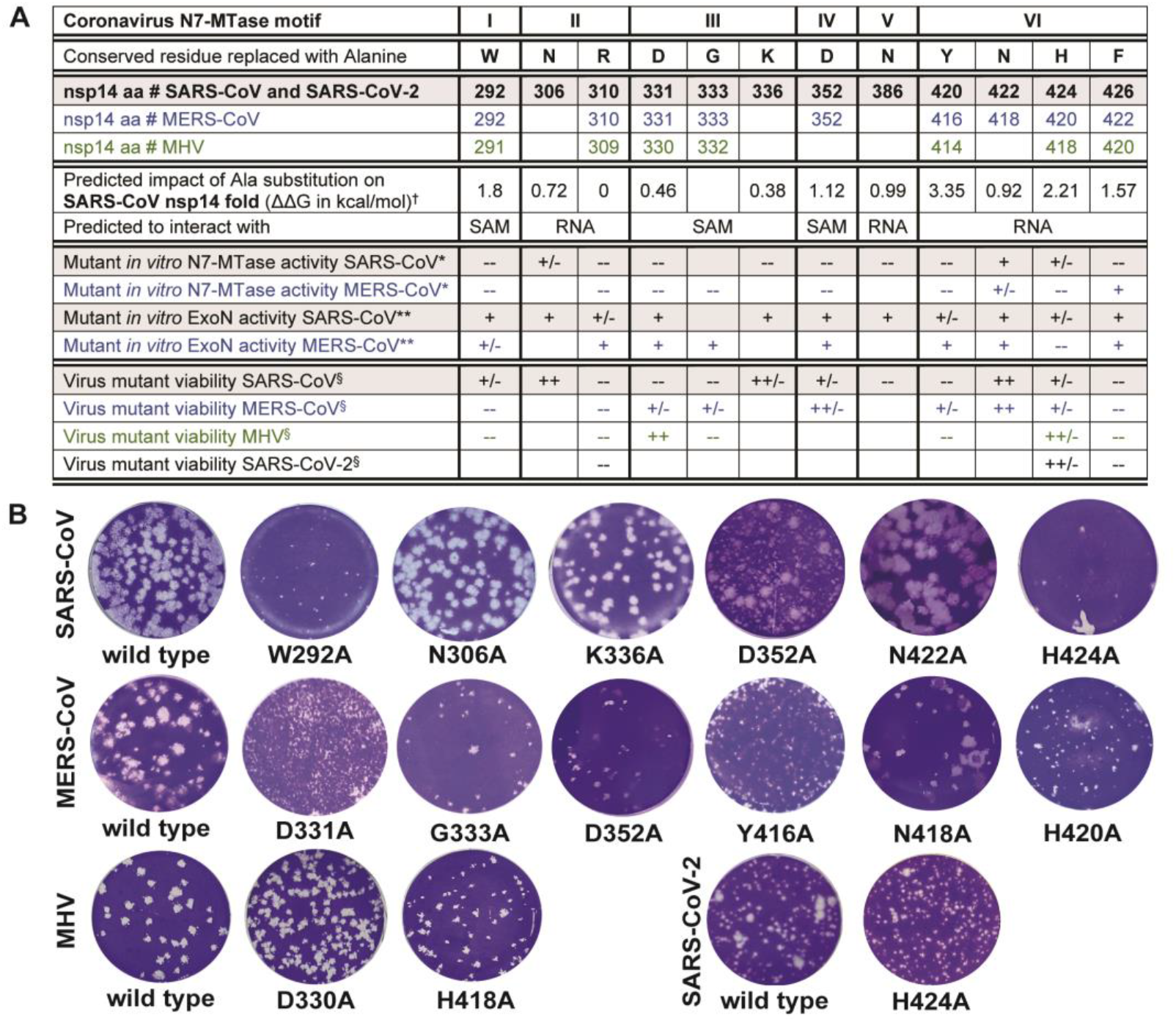
Virological characterization of betacoronavirus N7-MTase mutants. (A) Summary of results obtained from *in silico*, biochemical and virological studies of CoV mutants. ^†^ Values presented correspond to ΔΔG values from Table S2. * N7-MTase activity of each mutant was compared to the wild-type control enzyme and scored +, +/−, or -- when >50%, between 10 and 50%, or <10%, respectively. Values used here correspond to results obtained using the high enzyme concentration (Fig. 3B and 3D). ** ExoN activity of each mutant was evaluated relative to the wild-type control enzyme and scored +, +/−, or -- for equal, reduced, and abolished ExoN activity, respectively. ^§^ Mutant virus phenotypes, as deduced from plaque assays, were scored as: --, non-viable; +/−, severely crippled; ++/−, mildly crippled; ++, similar to the wild-type control. Empty cells indicate mutants that were not generated. (B) Plaque phenotype of the progeny of viable N7-MTase mutants. Plaque assays were performed using supernatants harvested from transfected cells at 3 (MERS-CoV in HuH7 cells, SARS-CoV and SARS-CoV-2 in Vero E6 cells) or 4 days (MHV in 17Cl1) post transfection.

In line with the biochemical data, the non-viable phenotype of six of the twelve SARS-CoV mutants (Fig. 5B) provided clear support for the importance of key residues in N7-MTase motifs II (R310), III (D331 and G333), V (N386), and VI (Y420 and F426). As anticipated, mutations in the canonical SAM binding motif III (DxGxPxG/A) completely abrogated SARS-CoV replication (Fig. 5A), apparently confirming the critical role of D331, which was postulated to be a key residue for methylation upon the discovery of the CoV N7-MTase (27). On the other hand, D331A was the only non-viable SARS-CoV mutant for which reversion to wild-type was occasionally observed, suggesting that a very low level of viral RNA synthesis remained possible in spite of this mutation (see also below).

Remarkably, SARS-CoV mutations N306A, K336A, and N422A in motifs II, III, and VI, respectively, were found to yield viruses with plaque phenotypes and progeny titers similar to those of the wild-type control (Fig. 5), despite the major impact of these mutations on *in vitro* N7-MTase activity (Fig. 3C). Likewise, the viable but severely crippled (small-plaque) virus phenotypes of motif-I mutant W292A and motif VI-mutant H424A were surprising (Fig. 5B), although for the latter the biochemical assays did reveal some activity when performed with an increased enzyme concentration (Fig. 3C and (35)). Interestingly, mutant D352A yielded a mixed-size plaque phenotype, suggesting rapid (pseudo)reversion in a minor fraction of this mutant’s progeny (Fig. 5B). For all six viable mutants, the presence of the original mutation in the viral progeny was confirmed by sequence analysis of the full-length nsp14-coding region of the viral genome. No other mutations were detected in this region of the genome. For non-viable mutants, transfected cells were incubated and monitored for 6 days and absence of viral activity was also confirmed by immunofluorescence microscopy with antibodies specific for double-stranded RNA and SARS-CoV nsp4.

In general, our data demonstrated the importance of the N7-MTase domain for SARS-CoV viability and confirmed the importance of the motifs and key residues identified using structural biology and biochemical approaches (summary presented in Fig. 5A). Nevertheless, for several mutants the data from different types of assays did not readily align, which prompted us to expand the reverse genetics efforts to other betacoronaviruses.

### Phenotypic differences between betacoronaviruses N7-MTase mutants suggest complex structure-function relationships

Even when targeting highly conserved viral functions, the introduction of equivalent mutations in closely related viruses can sometimes yield remarkably different mutant phenotypes. A recent example is the inactivation of the nsp14 ExoN, which is tolerated by MHV and SARS-CoV, but not by MERS-CoV and SARS-CoV-2, the latter virus having an nsp14 sequence that is 95% identical to that of SARS-CoV (37). To expand our understanding of the impact of N7-MTase mutagenesis, we engineered, launched, and analyzed a set of MERS-CoV and MHV mutants, using technical procedures similar to those described above for SARS-CoV (see Methods). In this case, the production of viable progeny was facilitated by co-culturing transfected BHK-21 cells with host cells appropriate for the amplification of MHV (17clone1 cells) or MERS-CoV (Huh7 cells). Again, each mutant was launched at least four times (from duplicate full-length cDNA clones) and the results are summarized in Fig. 5.

The mutations tested for MERS-CoV and MHV had a large predicted impact in our folding free energy analysis (Table S2 and Table S3) and/or yielded a non-viable or crippled phenotype in our SARS-CoV study (Fig. 5A). We evaluated whether these residues were equally critical for the replication of other betacoronaviruses. For clarity, in the text below we will refer to the conserved key residues of each motif instead of using nsp14 amino acid numbers, which are slightly different when comparing SARS-CoV, MERS-CoV and MHV (see Fig. 5A).

In contrast to the SARS-CoV result, the replacement of the W in SAM binding site motif I was lethal for both MERS-CoV and MHV. Strikingly, mutagenesis of the D and G in motif III (SAM binding site) yielded the opposite outcome: both were not tolerated in SARS-CoV, but resulted in crippled but viable or even wild type-like phenotypes for MERS-CoV and MHV, respectively (Fig. 5B). These results again indicated that CoV N7-MTase active site mutants can be (partially) viable, even in the absence of detectable *in vitro* enzymatic activity (Fig. 3D). Similar to our observations for SARS-CoV, the replacement of the D in motif IV and the N in motif VI had moderate or no impact, respectively, on the production of MERS-CoV progeny (Fig. 5). Replacement of the conserved H in motif VI (RNA binding site) consistently crippled replication across SARS-CoV, MERS-CoV, and MHV (Fig. 5B), while replacement of the conserved Y in the same motif was partially tolerated by MERS-CoV, but not by SARS-CoV and MHV.

Our betacoronavirus comparison identified only two N7-MTase mutations that consistently abrogated the replication of all three viruses tested: the R-to-A in motif II and the F-to-A in Motif VI, which both map to the putative RNA binding site. This was surprising in the case of MERS-CoV, given the fact that this mutation (F422A in MERS-CoV) allowed substantial N7-MTase activity in the *in vitro* assay (Fig. 3D). When SARS-CoV-2 emerged during the course of this study, the three mutations that produced a similar phenotype across SARS-CoV, MERS-CoV and MHV (R310A, H424A, and F426A; using SARS-CoV numbering) were also engineered for this newly discovered coronavirus. Again, the R310A and F426A replacements were found to fully abrogate virus replication, while H424A yielded a crippled phenotype in SARS-CoV-2 (Fig. 5).

## Discussion

Most viral MTases belong to the Rossmann-fold family (55, 59), a ubiquitous higher-order structure among dinucleotide-binding enzymes (55, 60). The CoV nsp14 N7-MTase was the first identified example of a non-Rossmann fold viral MTase (35, 36, 45), and the only one thus far for which some structural and functional information had been gathered. While some viral N7-MTase crystal structures have been resolved (35, 36, 61–63), their biochemical properties and signature sequences critical for RNA binding or enzymatic activity remain poorly defined compared to *e*.*g*. the 2′-O-MTases, an example of which is found in CoV nsp16 (reviewed in(6)). Likewise, the biological role and relevance of the CoV N7-MTase have not been explored in much detail. In recent studies and reviews, often related to SARS-CoV-2, the enzyme is widely assumed to secure the translation of CoV subgenomic mRNAs and genome, which obviously is a critical step for any positive-stranded RNA virus. However, direct biochemical evidence showing that CoV mRNAs indeed carry an N7-methylated cap at their 5’ end is still lacking. The presence of such a cap on CoV RNAs was first postulated following RNase T1 and T2 digestion studies with ^32^P-labeled MHV RNA, 40 years ago (64). Additional support came from immunoprecipitation experiments using a cap-specific monoclonal antibody (recognizing both the rare nucleoside 2,2,7-trimethylguanosine and 7-methylguanosine (m7G) cap structures) that brought down the mRNAs of equine torovirus (65), a distant CoV relative for which – perhaps strikingly – an N7-MTase domain still remains to be identified(45). The presence of enzymes required for capping in CoVs and many of their relatives (6, 17, 45, 47, 66) and the *in vitro* activity profile of recombinant CoV nsp14 (26, 27, 32, 33, 37, 38) lend additional credibility to CoV capping and cap methylation, but do not exclude the possibility that the CoV N7-MTase may target other substrates as well.

To enhance our overall understanding of nsp14 N7-MTase structure and function, also in the light of its emergence as an important drug target in the battle against SARS-CoV-2 (50, 67–69), we now revisited the SARS-CoV nsp14 X-ray structure to define the most likely residues involved in N7-MTase substrate binding and catalysis. Instead of a βαβ architecture (a seven-stranded β-sheet surrounded by six α-helices) and the canonical MTase motifs, the CoV N7-MTase incorporates twelve β-strands and five α-helices that form a five-stranded β-sheet core (36, 45). The overall nsp14 structure reveals two domains interconnected by a hinge that may confer the flexibility needed to orchestrate the different functions of the protein during CoV replication (36). Furthermore, the protein binds to nsp10, a critical co-factor for nsp14’s ExoN activity (34, 70). The conversion of a 5’-terminal GMP cap (GpppN) into a cap-0 structure (^7m^GpppN) involves multiple steps: stabilization of the RNA chain, SAM binding, methyl transfer to the N7 position of the cap, release of the methylated RNA substrate, and SAH release. Our structural analysis identified several residues with their side chains pointing towards the catalytic pocket, which could be classified as likely RNA- or SAM-binding motifs (Fig. 2B and 2C). Taking into account the amino acid sequence conservation between MHV, SARS-CoV, SARS-CoV-2, and MERS-CoV (Fig. 2A and alignment in Fig. S2), and the structures available to date (35, 36, 71, 72), we surmised these CoV N7-MTases to have an overall similar fold and structural organization. The impact of alanine substitutions of selected key residues in these motifs was then evaluated both *in vitro*, using SARS-CoV or MERS-CoV recombinant nsp14, and in the context of the viral replication cycle, by engineering the corresponding virus mutants in different betacoronaviruses.

Although the biochemical and virological data presented in this study clearly provide support for the predictions derived from our structural analysis, the overall interpretation of the data set undeniably is much more complex than anticipated (Fig. 5A). Replacement of conserved SARS-CoV and MERS-CoV N7-MTase residues largely or completely abrogated enzymatic activity *in vitro* (Fig. 3C and 3D), supporting their identification as key residues for the enzyme’s functionality when the protein is expressed alone (N7-MTase activity) or when tested in complex with nsp10 (ExoN activity). However, for several SARS-CoV and MERS-CoV mutations the data on enzymatic activity *in vitro* and virus mutant viability appeared to be at odds with each other (Fig. 5A). One possible interpretation is that (very) low levels of N7-MTase activity may still suffice to support viral replication in cell culture models. Alternatively, the *in vitro* N7-MTase assays may have suffered from technical complications, such as suboptimal or incorrect (mutant) N7-MTase domain folding. This could be different for nsp14 expressed in the context of the virus-infected cell and in the presence of its natural interaction partners, in particular other members of the viral replication and transcription complex. It is conceivable that the impact of nsp14 mutations on the fold and/or critical protein-protein or protein-RNA interactions of the N7-MTase domain could fluctuate between different assay systems. This might explain a stronger (*e*.*g*., MERS-CoV mutant F422A) or less dramatic effect in the virus-infected cell compared to what is observed in enzymatic assays (Fig. 5A). Mutations mapping to motif VI (hinge region) yielded inconsistent results in comparison to prior *in vitro* studies (26, 27, 32–35), which might be attributed (in part) to different *in vitro* assay conditions. Such technical explanations, however, do not apply when introducing equivalent substitutions in different betacoronaviruses and evaluating them in the context of the viral replication cycle. Also here apparent inconsistencies were observed in terms of the variable impact of certain mutations on the overall replication of virus mutants. The results obtained with mutations in motif III (the presumed SAM binding motif DxGxPxG/A) were a striking example: the viral phenotype for the D-to-A mutant (D331A in SARS-CoV and MERS-CoV, D330A in MHV) ranged from non-viable for SARS-CoV, via severely crippled for MERS-CoV to wild type-like for MHV (Fig. 5). SARS-CoV residue D331 was first identified as important for N7-MTase activity by the superimposition of nsp14 with cellular N7-MTase structures (27). However, a previous MHV study (43) had already documented that replacement of the corresponding residue D330 did not affect MHV replication, and pointed to G332 as a more important residue in motif III, which was confirmed in this study (Fig. 5). These results are consistent with the SARS-CoV nsp14 crystal structure showing that residue G333 in the DxG motif (G332 in MHV) is in direct contact with the SAM methyl donor (35), although apparently its replacement is not sufficient to render all betacoronaviruses non-viable. These results stress the importance to achieve a series of high-resolution structures of these different proteins in order to determine the subtle mechanistic differences.

The only other N7-MTase position probed by reverse genetics so far was the conserved tyrosine in motif VI (Fig. 2C; Y414 in MHV). This residue attracted attention by the intriguing serendipitous finding that its replacement with histidine did not affect replication of MHV strain A59 in cell culture, but strongly reduced replication and virulence in mice (41). Also, an Y414A substitution was tolerated in MHV-A59 (44, 58), but in our study Y414A prevented the recovery of infectious progeny for MHV strain JHM, which exhibits less robust RNA synthesis and overall replication than MHV-A59. The results for the corresponding SARS-CoV (non-viable) and MERS-CoV (crippled) mutants were also variable, adding to the complexity of the overall picture.

A substantial set of N7-MTase mutations was monitored for ‘side effects’ at the level of *in vitro* ExoN activity (Fig. 4), although for SARS-CoV and MHV these would unlikely explain a lack of viability as ExoN knock-out mutants for both these viruses are only mildly crippled (42, 58, 73). Strikingly, for MERS-CoV, which does not tolerate ExoN inactivation (37), two of the N7-MTase mutations (G333A in motif III and H420A in motif VI) abolished detectable ExoN activity *in vitro* (Fig. 4B), but still allowed a certain level of virus replication (small-plaque phenotype), an observation that clearly warrants further investigation. In more general terms, the ExoN biochemical assay (Fig. 4) suggested that the functional separation between the two enzyme domains may be less strict than previously concluded, as also recently hypothesized following an *in silico* and biochemical analysis using SARS-CoV-2 nsp14 ExoN domain (72, 73). Alternatively, structural variation may explain the discrepancies observed. The impact of SARS-CoV N7-MTase motif-VI mutations on ExoN activity was major, highlighting the peculiar structural organization of nsp14, in which part of the N7-MTase substrate-binding cavity maps to the hinge that connects the N7-MTase and ExoN domains (Fig. 1). For other N7-MTase motifs probed, the functional separation from ExoN was confirmed, as also deduced from previous studies (27, 33, 35, 38).

In our reverse genetics studies with four betacoronaviruses, a consistent phenotype was observed only for N7-MTase mutants carrying replacements of the conserved R in motif II (non-viable) and the conserved H and F in motif VI (crippled and non-viable, respectively). SARS-CoV residue R310 was previously reported to play a role in SAM binding (33), whereas F426 was proposed to entrench and stabilize the guanosine’s purine moiety in the proximity of SAM (35). Our analysis (Fig. 2) redefined both residues as part of putative RNA binding site motifs II and VI, respectively, and they were found to be essential for *in vitro* N7-MTase activity in SARS-CoV. Our results highlight the importance of the nsp14 N7-MTase for CoV replication, but the variable impact of the replacement of several conserved residues suggests a substantial degree of conformational or functional flexibility in the enzyme’s active site. Other factors, such as interactions of nsp14 with other replicase subunits, may also contribute to the observed phenotypic differences between equivalent N7-MTase mutants of different betacoronaviruses. Likewise, the translation of *in vitro* N7-MTase activity to virus viability is not straightforward and suggests complex structure-function relationships for the structurally unique CoV N7-MTase. Given both its essential role in CoV replication and its emerging status as a target for antiviral drug development efforts, it will be important to further expand the integrated biochemical and virological analysis to support the rational design of broad-spectrum inhibitors of the CoV N7-MTase.

## Materials and Methods

### Bioinformatics analysis

Forty-seven CoV nsp14 sequences were retrieved (a complete list is provided in Table S1) and aligned using MAFFT. Delineation of motif I to VI was done manually using Seaview and WebLogo (74, 75). Structure analysis (PDB: 5NFY; (36)), volume estimation, cavity determination and sequence conservation was plotted onto the structure using UCSF Chimera (76). Electrostatic surface calculations were done using APBS (77). Predicting the structural impact of mutations was done using the PoPMuSiC server (http://dezyme.com/en/software) (78). This program introduces single-site mutations into a protein’s structure and estimates the change in ΔΔGs values of such mutations. In the next step, all possible single-site mutations (4731 mutations) were sorted by their ΔΔGs, but only those in the conserved motifs in the vicinity of the catalytic pocket were used for further studies. PopMuSic predictions were cross-validated with SNAP2 to assess the impact of single amino acid substitutions on protein function (79).

### Recombinant protein expression and purification

Recombinant SARS- and MERS-CoV nsp10 and nsp14 were expressed in *E. coli* and purified as described previously (26), MERS-CoV-nsp14 (37, 49) and MERS-nsp10 (29, 80). Vectors for mutant nsp14 expression were generated by QuikChange site-directed mutagenesis using Accuzyme DNA polymerase (Bioline) and verified by sequence analysis. For each recombinant protein used, two batches were produced and tested in enzymatic assays.

### *In vitro* nsp14 N7-MTase activity assay

Reaction mixtures contained 50 or 200 nM of SARS-CoV or MERS-CoV recombinant nsp14, 7 nM GpppACCCC synthetic RNA substrate, 40 mM Tris-HCl (pH 8.0), 10 mM DTT, 5 mM MgCl_2_, 1.9 μM SAM, 0.1 μM 3H-SAM (Perkin Elmer). After a 30-min incubation at 30°C, the assay was stopped by addition of a 10-fold volume of ice-cold 100 μM S-adenosyl-homocysteine (SAH; Thermo Fisher). Samples were spotted on DEAE filter mats (PerkinElmer) and washed twice with 10 mM ammonium formate (Sigma-Aldrich) (pH 8.0), twice with MilliQ water, and once with absolute ethanol (Sigma-Aldrich) (26), and MTase activity was quantified using a Wallac scintillation counter. To determine relative enzyme activities, the incorporation measurements for mutant proteins were normalized to values obtained with wild-type nsp14. Samples were measured in triplicate in each experiment.

### *In vitro* nsp14 ExoN assay

Synthetic RNA substrate H4 (34) was radiolabeled at its 5’ end using T4 polynucleotide kinase (Epicentre) and [γ-^32^P]ATP (Perkin Elmer) and used as substrate in ExoN activity assays. To this end, recombinant SARS-CoV or MERS-CoV nsp14 and nsp10 were mixed in a 1:4 concentration ratio of nsp14:nsp10 as indicated in Fig. 4. The proteins were added to 500 nM radiolabeled substrate in reaction buffer (40 mM Tris-HCl (pH 7.5), 5 mM MgCl_2_, 1 mM DTT). The protein mix was left for 10 min at room temperature to allow the formation of the complex. Assays were performed at 37°C and stopped by addition of a 3x volume of loading buffer containing 96% formamide and 10 mM EDTA. Samples were analyzed on 7 M urea-containing 14% (wt/vol) polyacrylamide gels (acrylamide/bisacrylamide ratio, 19:1) buffered with 0.5xTris-taurine-EDTA and run at high voltage (1,600 V). Results were visualized by phosphorimaging using a Typhoon-9410 variable-mode scanner (GE Healthcare).

### Cell culture

Baby hamster kidney cells (BHK-21; ATCC CCL10), Vero E6 (ATCC; CCL-81), HuH7 cells and mouse 17 Cl1 cells were grown as described previously (19, 37, 81, 82). In order to amplify viral progeny and titrate recombinant CoVs by plaque assay, Vero E6 cells were used for SARS-CoV and SARS-CoV-2, HuH7 cells for MERS-CoV, and 17Cl1 cells for MHV. Cells were cultured in Eagle’s minimal essential medium (EMEM; Lonza) with 8% fetal calf serum (FCS; Bodinco) supplemented with 100 IU/ml of penicillin and 100 µg/ml of streptomycin (Sigma) and 2 mM L-Glutamine (PAA Laboratories). After infection, complete EMEM medium containing 2 % FCS was used.

### Viruses and reverse genetics

Mutations in the nsp14-coding region were engineered by two-step *en passant* recombineering in *E. coli* (83) using a bacterial artificial chromosome (BAC) vector with a full-length cDNA copy of a ß-CoV genome. Virus isolates used were MERS-CoV strain EMC/2012 (84, 85)), SARS-CoV Frankfurt-1 (86), MHV-JHM-IA (87), and SARS-CoV-2 BetaCoV/Wuhan/IVDC-HB-01/2019 (88). When designing mutations, additional translationally silent marker mutations was introduced near the site of mutagenesis, in order to analyze possible reversion and rule out potential contaminations with parental virus. For each mutant, two independent BAC clones were obtained, verified by sequencing of the full-length nsp14-coding region, and used for *in vitro* transcription (mMessage-mMachine T7 Kit; Ambion) and virus launching. Transfections with full-length RNA transcripts were performed as described before (37). Briefly, 5 µg RNA was electroporated into BHK-21 cells using an Amaxa nucleofector 2b (program A-031) and Nucleofection T solution kit (Lonza). Transfected BHK-21 cells were mixed in a 1:1 ratio with cells susceptible to CoV infection: Vero E6 cells (for SARS-CoV and SARS-CoV-2), HuH7 cells for MERS-CoV, or 17Cl1 cells (for MHV). Cell culture supernatants were collected when full cytopathic effect was observed, or at 6 days post transfection and progeny virus titers were determined by plaque assay (89). Viral replication was also monitored by immunofluorescence microscopy using antibodies recognizing double-stranded RNA (dsRNA;(90)) and non-structural or structural CoV proteins (37, 82, 91). To confirm the presence of the original mutations in viral progeny, supernatant from transfected cells was used to infect fresh cells, after which intracellular RNA was isolated with TriPure isolation reagent (Roche Applied Science). Next, the nsp14-coding region was amplified using standard RT-PCR methods and the purified amplicon was sequenced by Sanger sequencing. All work with live (recombinant) class-3 CoVs was done in a biosafety level 3 laboratory at Leiden University Medical Center.

## Data Availability

All study data are included in the article and SI Appendix.

## Acknowledgments

N.S.O. was supported by the Marie Skłodowska-Curie ETN European Training Network ‘ANTIVIRALS’ (EU Grant Agreement No. 642434). P.E.K. was the recipient of a scholarship from the Fundation “Méditerranée Infection”. This work was supported by the Fondation pour la Recherche Médicale (Aide aux Équipes) and the SCORE project (EU Horizon 2020 research and innovation program, grant agreement 101003627). We thank LUMC colleagues Tessa Nelemans for excellent technical support and Alexander Gorbalenya for scientific discussions and critical reading of the manuscript.

## Notes

### Competing Interest Statement

The authors have declared no competing interest.

### Summary of Updates

updated quality of Figure S2 in the support information file. Corrected some typos in the table S2 and Fig. 5A.

