## Supporting information for "Structure-function analysis of the nsp14 N7-guanine methyltransferase reveals an essential role in *Betacoronavirus* replication"

**This PDF file includes:**

Figures S1 and S2
Tables S1 to S3
SI References

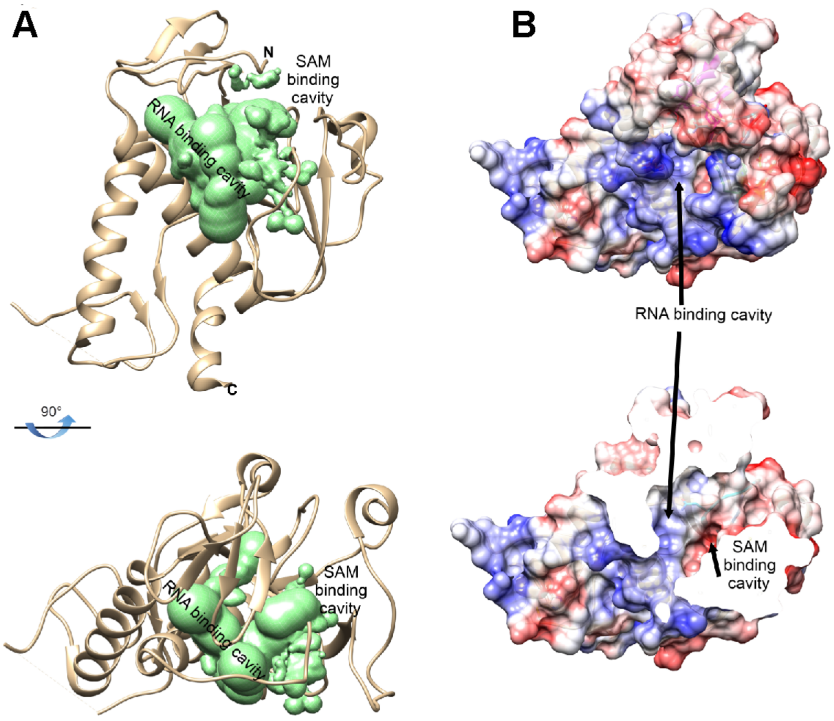


**Fig. S1.** Structural analysis of the catalytic site and hinge region of the SARS-CoV nsp14 N7-MTase domain. A) Determination of the volume of the enzyme's catalytic site (with the volume depicting the mold of the cavity shown in green). B) Electrostatic surface representation with the surface electrostatic potential calculated by APBS from -10 (red) to +10 (blue) kT/e.

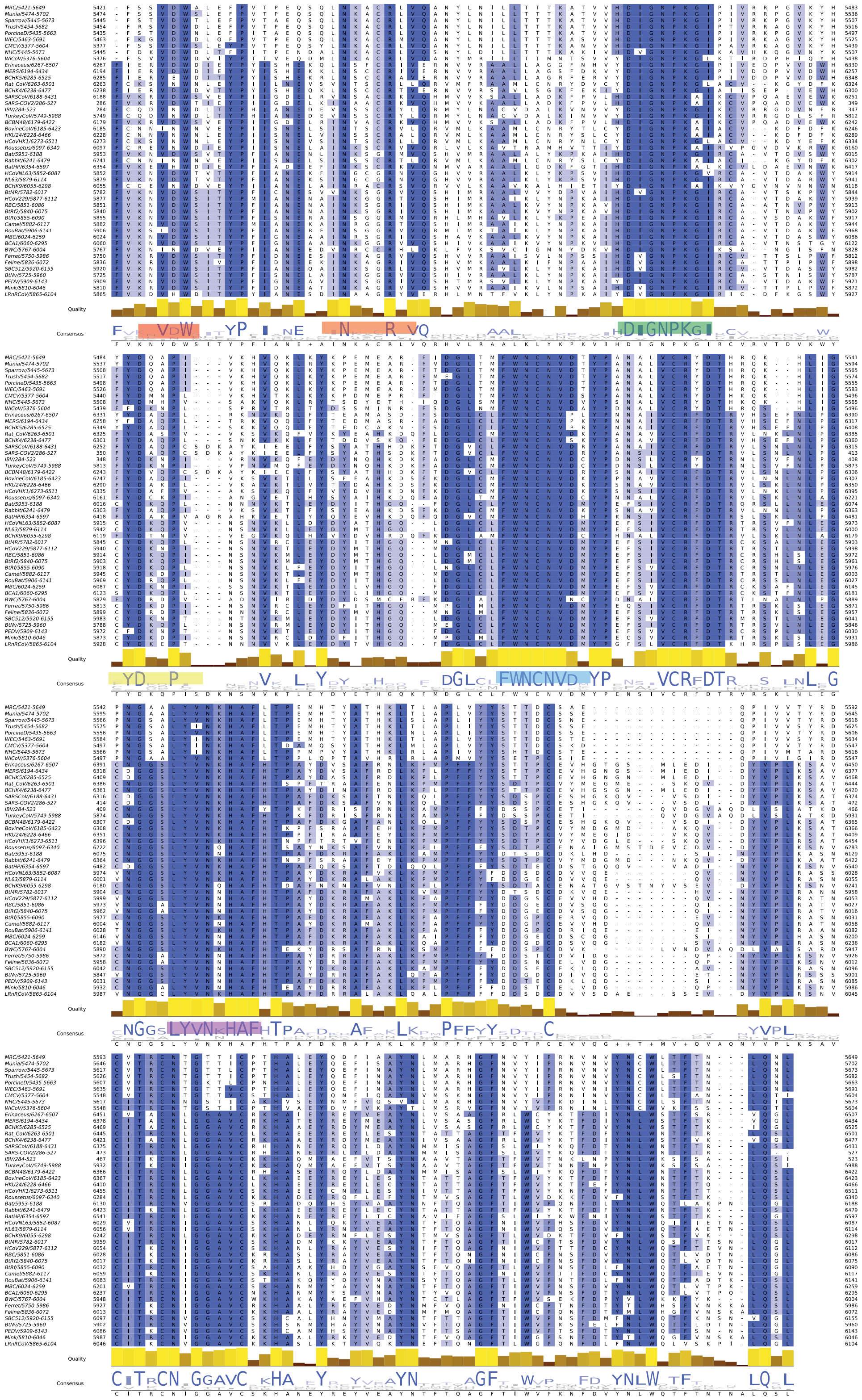

**Fig. S2.** Alignment of the nsp14 N7-MTase sequences from 47 selected coronaviruses (listed in Table S1). Highly conserved residues are boxed in dark blue (above 70% conservation), while partially conserved residues are displayed in lighter shades of blue. A consensus logo sequence is presented at the bottom of the alignment. The N7-MTase motifs I to VI are highlighted in red, orange, green, yellow, blue, and pink, using the same color scheme as in Fig. 2.

**Table S1.** List of coronaviruses and NCBI/Brenda-enzymes accession numbers used to extract nsp14 sequences for alignments (Fig. S2) and structural studies.

| Virus | Accession number |
|---|---|
| Alphacoronavirus BtMs-AlphaCoV/GS 2013 | A0A0U1WHG4 |
| Avian infectious bronchitis virus (IBV) | P0C6Y2 |
| Bat coronavirus 1A | YP_001718603.1 |
| Bat coronavirus BM48-31 | E0XIZ2 |
| Bat coronavirus CDPHE15/USA/200 6 | YP_008439224.1 |
| Bat coronavirus HKU4 | P0C6W3 |
| Bat coronavirus HKU5 | P0C6W4 |
| Bat coronavirus HKU9 | P0C6W5 |
| Bat Hp- betacoronavirus | A0A088DIE1 |
| Beluga whale coronavirus SW1 | YP_001876435.1 |
| Betacoronavirus Erinaceus | U5KNA9 |
| Betacoronavirus HKU24 | A0A0A7UXR0 |
| Bovine coronavirus | P0C6W8 |
| BtMr-AlphaCoV/ SAX2011 | A0A0U1UZC3 |
| BtNv-AlphaCoV/SC2013 | YP_009201729.1 |
| BtRf-AlphaCoV/HuB2013 | YP_009199789.1 |
| Camel alphacoronavirus | ALA50136.1 |
| Common moorhen coronavirus HKU21 | H9BR34 |
| Feline infectious peritonitis virus | AGZ84515.1 |
| Ferret coronavirus | YP_009256195.1 |
| Human coronavirus 229E (HCoV-229E) | P0C6X1 |
| Human coronavirus HKU1 (HCoV-HKU1) | P0C6X2 |
| Human coronavirus NL63 (HCoV-NL63) | P0C6X5 |
| Lucheng Rn rat coronavirus | YP_009336483.1 |
| Magpie-robin coronavirus HKU18 | H9BR07 |
| Middle East respiratory syndrome-related coronavirus | K9N7C7 |
| Miniopterus bat coronavirus HKU8 | YP_001718610.1 |
| Mink coronavirus strain WD1127 | YP_009019180 |
| Munia coronavirus HKU13-3514 | YP_002308505.1 |
| Murine coronavirus (strain A59) (MHV- A59) | P0C6X9 |
| Night heron coronavirus HKU19 | H9BR16 |
| Porcine deltacoronavirus | A0A140ESF0 |
| Porcine epidemic diarrhea virus | NP_839967 |
| Rabbit coronavirus HKU14 | H9AA60 |
| Rat coronavirus Parker | YP_009924380.1 |
| Rhinolophus bat coronavirus HKU2 | A8JNZ0 |
| Rousettus bat coronavirus | A0A1B3Q5W8 |
| Rousettus bat coronavirus HKU10 | AFU92103 |
| Scotophilus bat coronavirus 512 | YP_001351683 |
| Severe acute respiratory syndrome coronavirus (SARS-CoV) | P0C6X7 |
| Severe acute respiratory syndrome coronavirus-2 (SARS-CoV-2) | P0DTD1 |
| Sparrow coronavirus HKU17 | H9BQZ9 |
| Swine acute diarrhea syndrome related coronavirus BtRf2 | AVM80482.1 |
| Thrush coronavirus HKU12 | B6VDX7 |
| Turkey coronavirus | YP_001941187 |
| White-eye coronavirus HKU16 | YP_005352837.1 |
| Wigeon coronavirus HKU20 | H9BR24 |

**Table S2.** Projected impact on folding free energy by alanine substitutions of the identified core residues on the SARS-CoV N7-MTase structure, as calculated by PoPMusic (1).

| N7-MTase motif | Mutation | Secondary structure* | Solvent accessibility (%) | ΔΔG (kcal/mol) |
|---|---|---|---|---|
| | **wild-type** | - | - | 0 |
| **I** | **W292A** | T | 27.44 | 1.8 |
| **II** | **N306A** | H | 39.77 | 0.72 |
| | **R310A** | H | 58.37 | 0 |
| **III** | **D331A** | H | 4.20 | 0.46 |
| | **K336A** | H | 65.79 | 0.38 |
| **IV** | **D352A** | C | 14.21 | 1.12 |
| **V** | **N386A** | C | 32.59 | 0.99 |
| **VI** | **Y420A** | S | 86.78 | 3.35 |
| | **N422A** | S | 10.04 | 0.92 |
| | **H424A** | S | 43.12 | 2.21 |
| | **F426A** | C | 49.59 | 1.57 |

*classified as: T, turn; H, α-helix; C, random coil; and S, β-sheet.

**Table S3A.** Predicted functional impact of individual mutations of key conserved residues in motifs I-III of SARS-CoV, MERS-CoV, and MHV nsp14, predicted with SNAP2 (2).

| | **SARS-CoV** | **MERS-CoV** | **MHV** | **SARS-CoV** | **MERS-CoV** | **MHV** | **SARS-CoV** | **MERS-CoV** | **MHV** | **SARS-CoV** | **MERS-CoV** | **MHV** |
|---|---|---|---|---|---|---|---|---|---|---|---|---|
| **aa** | **W292** | **W292** | **W291** | **N306** | **N306** | **N305** | **R310** | **R310** | **R309** | **D331** | **D331** | **D330** |
| **A** | 72 | 66 | 78 | 51 | 26 * | 54 | 66 | 30 * | 63 | 75 | 67 | 78 |
| **R** | 82 | 81 | 88 | 75 | 32 * | 82 | | | | 91 | 80 | 90 |
| **N** | 80 | 63 | 86 | | | | 71 | 41 | 71 | 77 | 78 | 83 |
| **D** | 88 | 72 | 91 | 67 | 58 | 78 | 85 | 74 | 84 | | | |
| **C** | 59 | 62 | 69 | 48 | 22 * | 58 | 65 | 18 * | 63 | 73 | 65 | 76 |
| **Q** | 76 | 74 | 84 | 48 | 35 * | 65 | 60 | 32 * | 61 | 81 | 79 | 84 |
| **E** | 82 | 72 | 87 | 73 | 63 | 82 | 81 | 61 | 78 | 70 | 73 | 74 |
| **G** | 82 | 82 | 87 | 53 | 41 | 64 | 81 | 31 * | 79 | 79 | 84 | 88 |
| **H** | 78 | 62 | 86 | 50 | 36 * | 63 | 59 | 29 * | 60 | 87 | 84 | 89 |
| **I** | 72 | 69 | 79 | 71 | 35 * | 77 | 73 | 35 * | 71 | 88 | 85 | 89 |
| **L** | 73 | 75 | 82 | 72 | 35 * | 79 | 73 | 21 * | 70 | 90 | 81 | 91 |
| **K** | 85 | 84 | 90 | 63 | 59 | 80 | 41 | 20 * | 45 | 91 | 91 | 92 |
| **M** | 66 | 71 | 78 | 63 | 34 * | 73 | 66 | 29 * | 65 | 87 | 84 | 89 |
| **F** | 45 | 55 | 63 | 74 | 52 | 80 | 81 | 52 | 78 | 89 | 80 | 90 |
| **P** | 83 | 89 | 94 | 77 | 64 | 83 | 86 | 70 | 85 | 91 | 91 | 93 |
| **S** | 78 | 77 | 78 | 41 | 22 * | 58 | 70 | 32 * | 68 | 74 | 74 | 80 |
| **T** | 78 | 80 | 86 | 49 | 24 * | 64 | 69 | 32 * | 66 | 79 | 77 | 83 |
| **W** | | | | 81 | 70 | 86 | 87 | 66 | 85 | 92 | 88 | 93 |
| **Y** | 29 * | 50 | 52 | 70 | 54 | 77 | 79 | 53 | 77 | 90 | 83 | 91 |
| **V** | 71 | 60 | 78 | 67 | 26 * | 76 | 74 | 38 * | 72 | 87 | 83 | 88 |

SNAP2 scores are predicted with an expected accuracy above 70% confidence, except for scores labeled with *, which are predicted with an expected accuracy above 53% confidence. Positive values indicate a destabilizing effect, while negative values indicate a neutral effect.

**Table S3B.** Predicted functional impact of individual mutations of key conserved residues in motifs III-VI of SARS-CoV, MERS-CoV, and MHV nsp14, predicted with SNAP2 (2).

| | **SARS-CoV** | **MERS-CoV** | **MHV** | **SARS-CoV** | **MERS-CoV** | **MHV** | **SARS-CoV** | **MERS-CoV** | **MHV** | **SARS-CoV** | **MERS-CoV** | **MHV** |
|---|---|---|---|---|---|---|---|---|---|---|---|---|
| **aa** | **K336** | **K336** | **K335** | **D352** | **D352** | **D349** | **N386** | **N382** | **N380** | **Y420** | **Y416** | **Y414** |
| **A** | 31 * | 20 * | 39 * | 34 * | 63 | 73 | 60 | 45 | 56 | 71 | 66 | 74 |
| **R** | -17 * | -20 * | -1 * | 45 | 85 | 87 | 78 | 73 | 77 | 89 | 85 | 91 |
| **N** | 28 * | 26 * | 44 | 22 * | 73 | 69 | | | | 86 | 71 | 88 |
| **D** | 69 | 63 | 76 | | | | 62 | 56 | 63 | 90 | 73 | 92 |
| **C** | 22 * | 7 * | -16 * | 27 * | 57 | 69 | 60 | 15 | 57 | 61 | 54 | 64 |
| **Q** | 16 * | 6 * | 29 * | 43 | 74 | 73 | 47 | 52 | 64 | 82 | 77 | 86 |
| **E** | 52 | 45 | 60 | 26 * | 67 | 53 | 75 | 70 | 74 | 87 | 82 | 89 |
| **G** | 52 | 41 | 60 | 49 | 77 | 78 | 51 | 55 | 49 | 87 | 81 | 88 |
| **H** | 7 * | -6 * | 19 * | 21 * | 74 | 80 | 63 | 51 | 63 | 59 | 70 | 78 |
| **I** | -15 * | -70 | -40 * | 59 | 78 | 86 | 77 | 53 | 76 | 69 | 62 | 70 |
| **L** | 22 * | 8 * | 33 * | 61 | 80 | 88 | 80 | 49 | 78 | 74 | 70 | 74 |
| **K** | | | | 48 | 87 | 88 | 76 | 71 | 76 | 90 | 86 | 91 |
| **M** | 10 * | -39 * | 18 * | 55 | 78 | 84 | 74 | 52 | 73 | 78 | 72 | 79 |
| **F** | 52 | 41 | 55 | 23 * | 71 | 87 | 81 | 67 | 80 | 42 | 47 | 40 |
| **P** | 62 | 30 * | 69 | 72 | 89 | 89 | 71 | 76 | 82 | 88 | 91 | 94 |
| **S** | 22 * | 13 * | 36 * | 29 * | 65 | 65 | 44 | 31 | 42 | 85 | 69 | 86 |
| **T** | 26 * | 13 * | 39 * | 34 * | 69 | 69 | 51 | 30 * | 33 * | 86 | 79 | 88 |
| **W** | 67 | 56 | 67 | 66 | 81 | 90 | 87 | 78 * | 86 | 68 | 71 | 71 |
| **Y** | 48 | 35 * | 51 | 32 * | 76 | 86 | 79 | 38 * | 78 | | | |
| **V** | 22 * | 8 * | 32 * | 53 | 76 | 83 | 75 | 41 | 75 | 68 | 49 | 70 |

SNAP2 scores are predicted with an expected accuracy above 70% confidence, except for scores labeled with *, which are predicted with an expected accuracy above 53% confidence. Positive values indicate a destabilizing effect, while negative values indicate a neutral effect.

**Table S3C.** Predicted functional impact of individual mutations of key conserved residues in motif VI of SARS-CoV, MERS-CoV, and MHV nsp14, predicted with SNAP2 (2).

| | SARS-CoV | MERS-CoV | MHV | SARS-CoV | MERS-CoV | MHV | SARS-CoV | MERS-CoV | MHV |
|---|---|---|---|---|---|---|---|---|---|
| **aa** | **N422** | **N418** | **N416** | **H424** | **H420** | **H418** | **F426** | **F422** | **F420** |
| **A** | 66 | 33 * | 69 | 63 | 62 | 64 | 56 | 59 | 67 |
| **R** | 81 | 72 | 84 | 74 | 26 * | 35 * | 83 | 81 | 85 |
| **N** | | | | 63 | 70 | 65 | 82 | 79 | 83 |
| **D** | 64 | 65 | 79 | 84 | 86 * | 85 | 89 | 88 | 90 |
| **C** | 61 | 41 | 66 | 64 | 20 | 64 | 48 * | 39 * | 50 |
| **Q** | 64 | 4 * | 70 | 65 | 69 | 67 | 78 | 72 | 80 |
| **E** | 80 | 71 | 83 | 79 | 81 | 80 | 84 | 72 | 85 |
| **G** | 64 | 20 * | 64 | 75 | 66 | 76 | 81 | 79 | 83 |
| **H** | 60 | 49 | 67 | | | | 59 | 76 | 79 |
| **I** | 79 | 71 | 83 | 76 | 69 | 76 | 58 * | 10 * | 60 |
| **L** | 81 | 72 | 85 | 76 | 41 | 75 | 63 | 50 | 59 |
| **K** | 79 | 69 | 81 | 81 | 81 | 82 | 85 | 82 | 87 |
| **M** | 76 | 63 | 80 | 72 | 65 | 72 | 60 | 49 | 47 |
| **F** | 69 | 74 | 85 | 64 | 75 | 76 | | | |
| **P** | 84 | 76 | 87 | 90 | 90 | 90 | 89 | 87 | 91 |
| **S** | 52 | -3 * | 59 | 70 | 71 | 72 | 79 | 75 | 80 |
| **T** | 59 | 42 | 65 | 64 | 72 | 76 | 78 | 60 | 80 |
| **W** | 88 | 81 | 90 | 81 | 82 | 82 | 69 | 69 | 67 |
| **Y** | 79 | 71 | 83 | 65 | 70 | 67 | 49 | 49 | 49 |
| **V** | 78 | 65 | 81 | 71 | 54 | 72 | 62 * | 37 * | 65 |

SNAP2 scores are predicted with an expected accuracy above 70% confidence, except for scores labeled with *, which are predicted with an expected accuracy above 53% confidence. Positive values indicate a destabilizing effect, while negative values indicate a neutral effect.
